# Cellular Protrusions Engage Viral Infection Enhancing EF-C Peptide Nanofibrils

**DOI:** 10.1101/2020.10.01.321810

**Authors:** Desiree Schütz, Sascha Rode, Clarissa Read, Janis A. Müller, Bernhard Glocker, Konstantin Sparrer, Oliver Fackler, Paul Walther, Jan Münch

## Abstract

Self-assembling peptide nanofibrils (PNF) have gained increasing attention as versatile molecules in material science and biomedicine. One important application of PNF is to enhance retroviral gene transfer, a technology that has been central to the development of gene therapy. The best-investigated and commercially available PNF is derived from a 12-mer peptide termed EF-C. The mechanism of transduction enhancement depends on the polycationic surface of EF-C PNF, which binds to the negatively charged membranes of viruses and cells thereby overcoming electrostatic repulsion and increasing virion attachment and fusion. Assuming an even distribution of charges at the surfaces of virions and cells would result in an evenly distributed interaction of the virions with the cell surface. However, we here report that PNF do not randomly bind at the cell surface but are actively engaged by cellular protrusions. Chemical suppression of protrusion formation in cell lines and primary CD4+ T cells greatly reduced fibril binding and hence virion binding. Thus, the mechanism of PNF-mediated viral transduction enhancement involves active engagement of virus-loaded fibrils by cellular protrusions.

## Introduction

Retroviral gene transfer is the method of choice for the stable introduction of genetic material into the cellular genome and is widely used in basic and translational research including gene therapy approaches^[1, 2]^. Efficient gene transfer is, however, often limited because of low transduction efficiencies which is mainly due to low vector titers and electrostatic repulsions between negatively charged viral and cellular membranes^[3]^. EF-C peptide nanofibrils (PNF, Protransduzin®) represent a novel and versatile tool to overcome these limitations^[4]^. These fibrils enable a convenient concentration of retroviral vector particles by low speed centrifugation and potently enhance virion attachment to target cells, which results in increased transduction efficiencies^[4]^. EF-C is a 12-mer peptide (QCKIKQIINMWQ) derived from the HIV-1 glycoprotein gp120 that instantaneously assembles into amyloid-like fibrils when dissolved in polar solvents such as water or cell culture media^[4]^. The fibrils have a diameter of ~ 3 nm, are hundreds of nanometers long, show a positive ζ potentials at neutral pH and are thus polycationic^[4,5]^.

EF-C fibrils are at least as active in increasing transduction efficiencies as RetroNectin^[4]^. RetroNectin is the gold standard transduction enhancer for retroviral/lentiviral gene transfer to hematopoietic cells which holds great promise in clinical application, having been used for over 40 clinical trials. This fibronectin derivative is typically coated on the bottom of cell culture dishes to capture viral particles and bringing them in close proximity to target cells^[4,6]^. However, the RetroNectin-coating procedure is cumbersome with time consuming liquid handling steps. In contrast, EF-C PNF offer a significantly simplified option since they are directly added with the viral particles to the target cells. Thus, EF-C PNF are currently evaluated as transduction enhancer to accelerate production of genetically reprogrammed immune cells, such as T-lymphocytes with chimeric antigen receptors in CAR-T immunotherapy^[7]^. Specifically, in connection with mini bioreactors, EF-C PNF may enable an *en gros* production of CAR-T cells by allowing transduction and expansion of cells in one reaction container^[7]^.

Transduction enhancement by EF-C PNF mediated through electrostatic interactions between the positively charged fibrils and negatively charged viral and cellular membranes^[8,9]^. This overcomes the charge repulsions of the membranes and results in increased rates of virion attachment and fusion with target cells^[5]^. The mechanism is supported by data showing that abrogating the cationic properties of the fibrils through anionic polymers diminishes their ability to enhance infection^[10]^. This is supported by the finding that the broad activity of EF-C PNF on retroviral vectors is largely independent of the type of incorporated glycoproteins. It is noteworthy that EF-C PNF increase retroviral transduction more efficiently than other commercially available polycations, e.g. soluble dextran or polybrene^[4]^. Thus, the polycationic charge alone does not explain the superior performance of EF-C PNF over other transduction enhancers, suggesting that the fibrillar structure may also contribute to transduction enhancement.

## Results

To better understand the transduction enhancing mechanism of the EF-C PNF, we here explored how fibrils interact with the cell surface. For this, we directly visualized the interaction of EF-C PNF with the surface of HeLa cells applying electron microscopy. Cells were grown on glass cover slips and incubated for 1 hour with buffer or 5 μg/ml of EF-C PNF which represents a standard concentration for efficient viral transduction enhancement ^[4,11]^. After sample fixation and preparation, cells were imaged by scanning electron microscopy (SEM). EF-C PNF were clearly detectable as fibrillar mesh (Figure 1B-D) which was absent in the untreated cells (Figure 1A). We did not observe differences in the size of treated and untreated cells. The fibrils were easily detectable at higher magnifications (Figure 1D) and are characterized by bright and filigree structures on the cell surface (see red arrows). We assumed that if electrostatic interactions are the only driving force, the fibrils will be randomly distributed on the cell surface. To our surprise, however, the fibrils clustered to cellular protrusions, i.e. filopodia, which seemed to actively engage or capture the fibrils, (Figure 1B-D). Time-lapse fluorescence microscopy indeed revealed that HeLa cells capture fluorescent EF-C PNF^[11]^ via filopodia and pull them within minutes to the cell surface (Movie S1 and S2).

**Figure 1:**
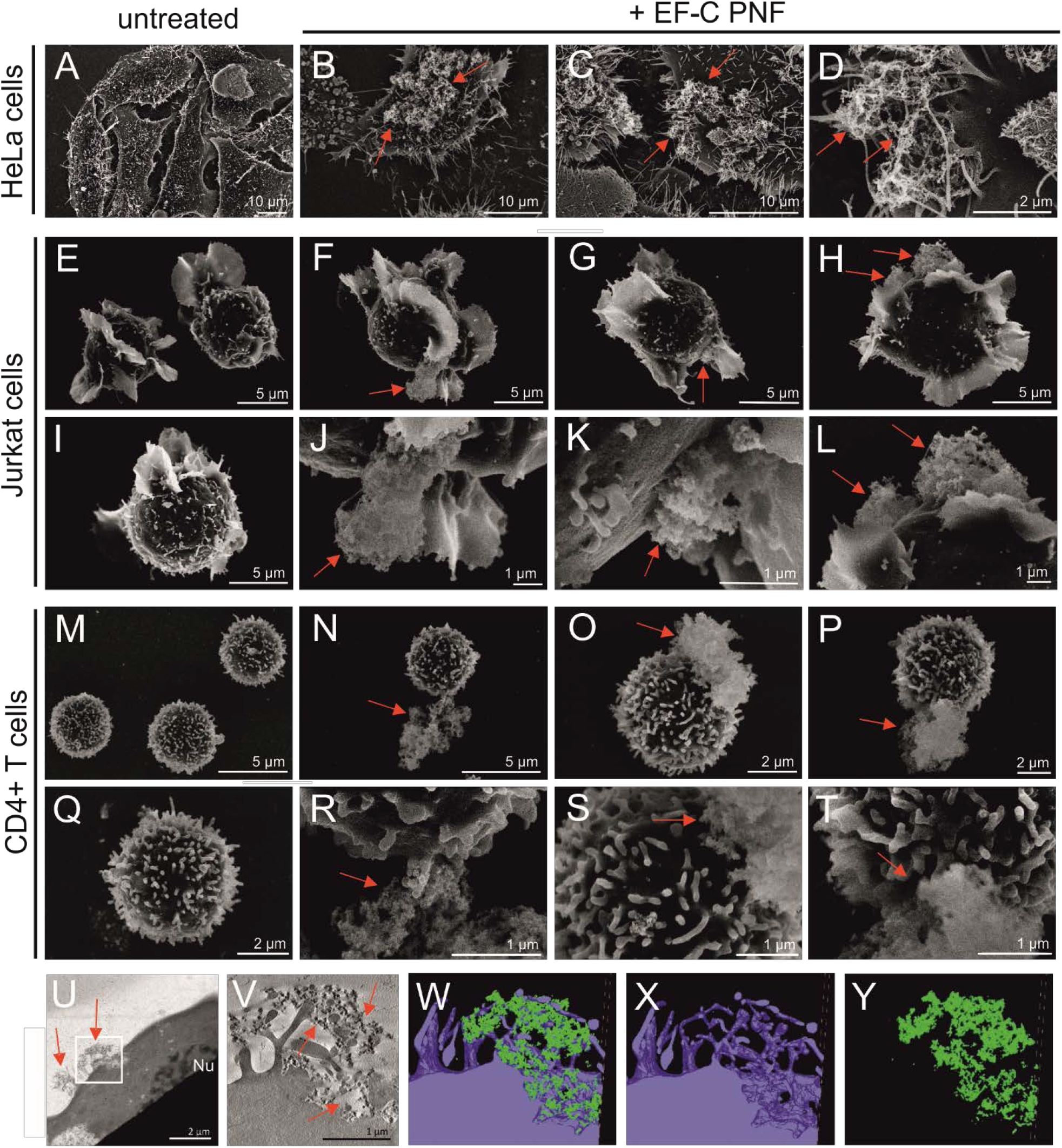
Fibrils bind to cellular protrusions. Analysis of EF-C binding to HeLa (A-D), Jurkat (E-L) and CD4 - T cells (M-T) visualized by scanning electron microscopy (SEM). Cells were incubated for 1 h at 37°C with 5μg/ml EF-C followed by fixation. Untreated cells are represented in A, E, I, M and Q. J-L and R-T are magnified sections of F-H and N-O respectively. Red arrows indicate EF-C PNF on cell surface. U-Y: HeLa cells were incubated with 1mg/ml EF-C PNF (FRET labelled) for 24h and analyzed by scanning transmission electron microscopy (STEM) (U). 3D datasets were obtained by tomography (V-Y). Virtual section of the tomogram of the indicated area in U (V). Nu: Nucleus. W-Y: 3D visualization of the tomogram from the same area. Merged (W) or plasma membrane (blue, X) and EF-C PNF (green, Y) alone.

We next investigated the interaction of EF-C fibrils with T cells, the target cells for e.g. CAR T cell therapy^[12]^. SEM analysis of immortalized Jurkat T cells (Figure 1 E-L) and primary CD4+ T cells (Figure 1 M-T) showed that after 1 hour of incubation the fibrils were as well specifically engaged by cellular protrusions. Due to the different cell morphologies of T cells compared to HeLa cells, this were mainly lamellipodia (Jurkat T cells) and, microvilli (primary CD4+ T cells). For better spatial evaluation of the interaction with fibrils and protrusion with higher resolution, we finally performed scanning transmission electron microscopy (STEM) tomography of HeLa cells incubated with EF-C PNF for 24 hours. As shown in Figure 1U and 1V,treatment of cells with EF-C PNF causes invaginations of the plasma membrane. The 3D visualization of this invagination clearly shows a complex mesh of cellular protrusions (Fig. 1X) engaging a cluster of fibrils (Fig. 1W and 1Y). Thus, electron microscopy and time-lapse fluorescence microscopy evaluation of the early events of the EF-C PNF interaction with cellular surfaces demonstrated that fibrils not just randomly bind to the cellular surface but are actively engaged by cellular protrusions and suggests that fibril-cell interaction causes formation of plasma membrane invaginations.

To further study the interaction of PNF with cellular protrusions, we took advantage of dynasore, a dynamin inhibitor which suppresses filopodia and lamellipodia formation, as previously described^[13–15]^. We first determined possible cytotoxic effects of dynasore for HeLa and Jurkat T cells but did not observe a reduction in metabolic activity at concentrations of up to 400 μM (Figure S1). Next, confocal microscopy was applied to visualize the effect of the dynamin inhibitor on cell morphology and filopodia formation. As shown in Figures S2A and S2B, a dynasore concentration of 200 μM resulted in a significant reduction of filopodia in both, HeLa and Jurkat T cells.

To quantitatively assess the effect of dynasore on the interaction of PNF with cells, HeLa cells were treated with mock or increasing concentrations of dynasore followed by incubation with fluorescent EF-C PNF^[11]^. Flow cytometry analyses revealed a time-dependent increase of EF-C positive cells from 15 % at 10 min to 70 % at 60 min (Figure S2C). Pretreatment of cells with dynasore concentrations of ≥ 200 μM effectively prevented fibril interaction (Figure S2C). Thus, HeLa cells treated with dynasore do not form protrusions and have a greatly reduced ability to interact with EF-C PNF.

Next, HeLa, Jurkat and CD4+ T cells were exposed to 200 μM dynasore, incubated with fluorescent fibrils over time and analyzed via flow cytometry. Again, time-dependent increase of PNF binding to HeLa cells was observed (Figure 2A). In contrast, 43 % of Jurkat T cells (Figure 2B) and 44 % of the primary CD4+ T cells (Figure 2C) in suspension culture were already associated with fibrils after 10 min, and these values only marginally increased after 60 min to ~ 60 % and ~ 52 % for Jurkat and CD4- T cells, respectively (Figure 2B and C). Dynasore treatment again abrogated fibril binding and resulted in average in ~10 % positive Jurkat and ~ 20 % positive CD4+ T cells. Taken together, EF-C PNF seem to interact with suspension cells with faster kinetics than with adherent HeLa cells and pharmacological inhibition of cellular protrusions greatly reduces the ability of adherent and suspension cells to capture EF-C PNF.

**Figure 2:**
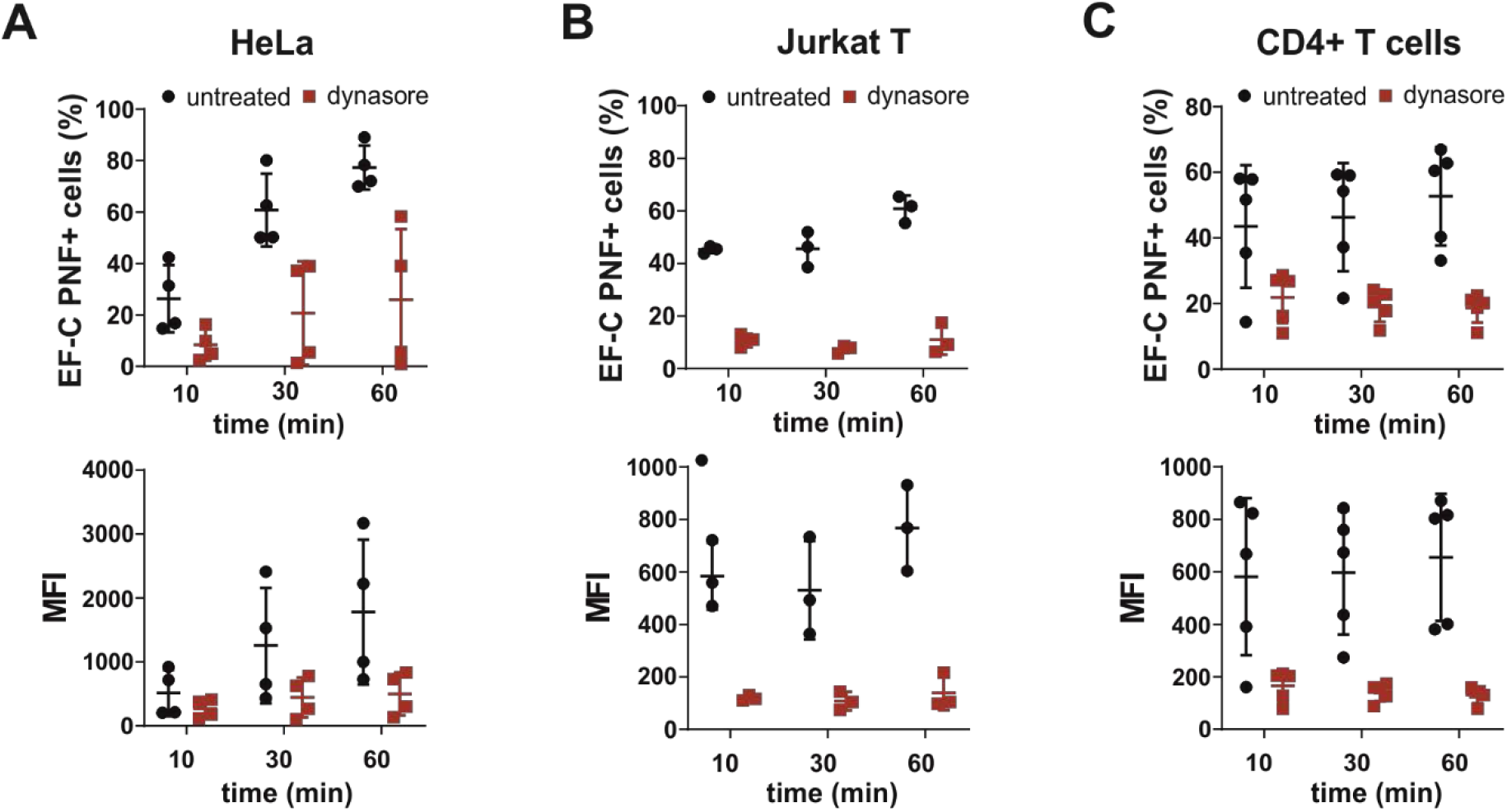
Cellular protrusions contribute to cellular EF-C PNF binding. ATTO495 labeled EF-C PNF was added to HeLa (A), Jurkat (B) and CD4 T (C) cells after a 45 min treatment with 200 μM dynasore. After indicated timepoints cells were washed, and the percentage of EF-C PNF positive cells and mean ATTO495 fluorescence intensity (MFI) were assessed by flow cytometry. Shown are average values (± SD) of triplicate measurements from three to five independent experiments.

Having shown that suppression of cellular protrusions abrogates PNF binding, we wondered whether induction of protrusions increases the interaction of fibrils with the cell surface. Blebs or membrane protrusions occur in a variety of cells upon physiological stress or during viral infections^[16,17]^. First, we tested whether transduction of HeLa cells with lentiviral vectors expressing shRNAs induces bleb formation. In fact, transduction resulted in an increased number of cells with blebs compared to non-transduced cultures (Figure S3A, S3B). We next determined the interaction of fluorescent EF-C PNF with transduced versus non-transduced cells via flow cytometry, and found that transduced cells exhibited increased mean fluorescence intensities (Figure S3C, S3E) and an increased ability to bind EF-C PNF (Figure S3D). Thus, induction of cellular protrusions results in enhanced EF-C PNF binding.

EF-C PNF bind viral particles and increase rates of viral attachment and their fusion to target cells^[4,11]^. Thus, cells lacking protrusions should also have a reduced capability to bind virus-loaded PNF resulting in reduced virus fusion. To test this hypothesis, HIV-1 fusion with dynasore-treated TZM-bl cells (a HeLa-derived cell line) was analyzed using the FACS-based HIV-1 Vpr BLAM fusion assay^[18]^. However, we observed that dynasore concentrations required to suppress filopodia formation (≥ 100 μM) effectively inhibit viral fusion already in the absence of fibrils, showing that this assay is not suitable to determine the relevance of protrusions for PNF-mediated transduction enhancement (Figure S4). Thus, we directly quantified rates of virion attachment to cells by confocal microscopy. For this, PBS or dynasore treated HeLa cells were exposed for 1 h to viral particles alone or in the presence of EF-PNF. Thereafter, cells were washed and viral capsids were stained with an anti-p24-antibody, imaged by confocal microscopy (Figure 3A) and viral particles were counted (Figure 3B). Dynasore alone did not significantly affect virion binding to the cell surface. In the presence of EF-C PNF, viral attachment was enhanced by a factor of 4.3, which was greatly reduced in dynasore-treated cells. These data show that protrusions also capture virus-loaded EF-C PNF.

**Figure 3:**
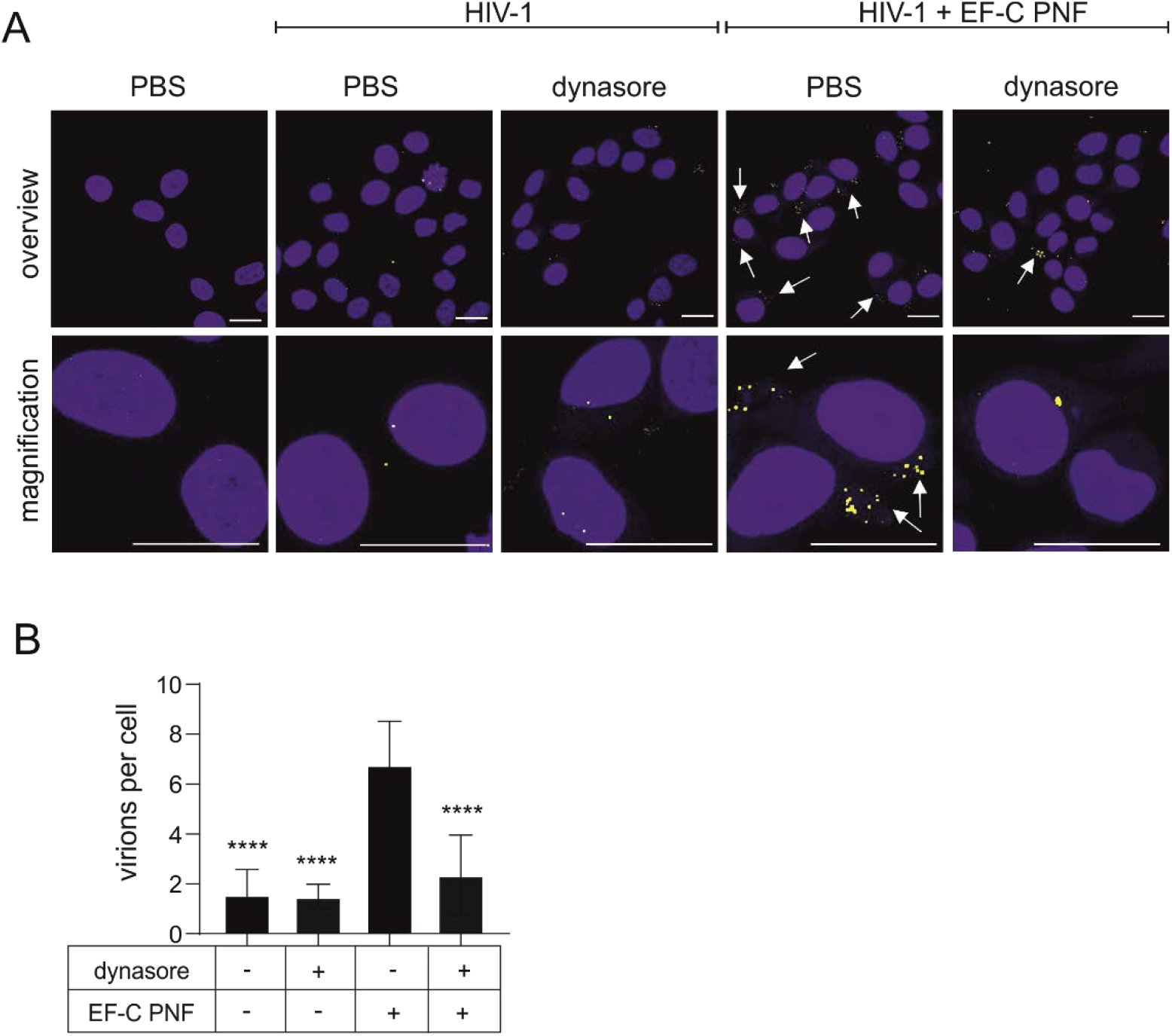
Dynasore abrogates binding of virus-loaded fibrils to the cell surface. A: Confocal microscopy of HeLa cells treated for 1h with HIV-1 or HIV-1 + EF-C PNF in presence or absence 200 μM Dynasore [blue = DAPI staining; yellow = p24 staining; white arrow = EF-C PNF]. Scale bars represent 20 μm. B: Quantification of viral particles per cell was determined by counting nuclei and yellow foci of six images for each sample. Shown are average values (± SD) of sextuplicate measurements from one experiment. Statistical analysis using ordinary one-way ANOVA shows significant differences between all samples and EF-C PNF only.

## Discussion

EF-C PNF are a versatile tool to increase retroviral transduction rates *in vitro* but also *ex vivo*, e.g. in CAR T cell therapy. It is therefore of great interest to clarify how these nanomaterials interact with relevant target cells. Our finding that cellular protrusions engage EF-C PNF came as a surprise, because we and others assumed that electrostatic interactions are the only driving force that mediates binding of the positively charged fibrils to the negatively charged cell surface ^[4,10,19]^. Thus, besides electrostatic interactions, the suprastructure of fibril aggregates itself seems to be an important determinant for cell binding and activity. Electron microscopy and time-lapse fluorescence microscopy analysis revealed that in particular filopodia actively engage EF-C PNF and pull them towards the cell body, which may explain the increased viral attachment rates in the presence of PNF. The virions are brought in close proximity to the cell membrane, where the viral entry process can be initiated through engagement of specific cellular receptors resulting in direct fusion with the cellular membrane or endocytotic uptake. Notably, STEM tomography indicated that engagement of fibril aggregates by cellular protrusions also induces invaginations of the plasma membrane, thereby forming macropinosomes, which may also serve as an entry route for many viruses or viral vectors ^[20]^. Thus, we propose that fibrils not only bind but also deliver viral particles to the cell body thereby facilitating infection. This may explain why PNF have a superior transduction enhancing activity as compared to soluble polycations such as polybrene or DEAE-dextran ^[4,21,22]^ because the latter do not form a fibrillar superstructure and are presumably not captured by protrusions.

Cellular protrusions are highly dynamic structures involved in fundamental processes, including cell migration and invasion, but also environmental sensing and cellular uptake mechanisms ^[23,24]^. In lamellipodia and filopodia actin polymerization drives forward protrusion of the plasma membrane^[25]^. We show that pharmacological inhibition of actin polymerization reduced the interaction of EF-C PNF with adherent and suspension cells, and, vice versa, that stress-induced induction of protrusions resulted in increased fibril binding. Interestingly, we also found that dynasore treatment did not completely abrogate fibril binding, because ~ 10 to 20 % of the cells always retained fibril binding activity. These findings suggest that either dynasore treatment did not completely abrogate fibril formation, or that this background binding of fibrils to cells is mediated by only electrostatic interactions independent of protrusions.

Cellular attachment of viruses is the rate limiting step in viral infection and vector transduction. It is generally accepted that EF-C fibrils enhance viral transduction by increasing the rates of virion attachment to and infection of target cells ^[4]^. Thus, we also aimed at analyzing the role of cellular protrusions in the presence of infectious virus. However, pharmacological inhibition of protrusions by dynasore affected viral infection even in the absence of fibrils (data not shown and ^[26,27]^). Thus, we sought to analyze the role of protrusions in a virion fusion assay, but again observed that dynasore affects viral fusion with the membrane precluding a meaningful analysis of the role of the fibrils in this process. We therefore directly quantified the number of virions that bound to dynasore-treated and untreated cells, in the absence or presence of fibrils. As expected from previous studies ^[4]^, fibrils increased viral attachment rates. Under these conditions, dynasore treatment did not affect virion binding to cells, but prevented attachment of virions captured by fibrils. In conclusion, protrusions not only bind fibrils but also virus-loaded fibrils, arguing that this process is mainly responsible for infection/transduction enhancement.

Conclusively, we here show that cellular protrusions actively engage transduction enhancing EF-C PNF and brings them into close proximity of the cell body which may result in increased attachment, fusion and infection rates. Fibril binding initiates the formation of invaginations of the cellular membrane, suggesting that fibrils may also get internalized. Follow-up studies to clarify the intracellular fate of the fibrils, e.g. whether they are degraded or stored as fibrils, are highly relevant for gene therapy studies in which transduced and fibril-associated or fibril-containing cells are reintroduced into the patient’s bloodstream.

## Experimental Section

A detailed description of material and methods is available in the Supporting Information Section.

## Supporting Information

Supporting Information is available from the Wiley Online Library or from the author.

## Acknowledgement

This work was supported by grants from the DFG (Collaborative Research Centre 1279 “Exploiting the Human Peptidome for Novel Antimicrobial and Anticancer Agents”) to J.M. and P.W., the Volkswagenstiftung to J.M., and the Leibniz Association to J.M.

