## supplement for "Cellular Protrusions Engage Viral Infection Enhancing EF-C Peptide Nanofibrils"

### Experimental Section

*Cell culture:* Adherent TZM-bl reporter cells, herein referred as HeLa cells, (NIH Aids Reagent) HEK 293T cells (ATCC, CRL-11268) and HeLa cells stably expressing LifeAct.GFP (kindly provided by Oliver Fackler, Department of Infectious Diseases, Virology, Heidelberg) were cultured in DMEM with 120 µg/mL penicillin, 120 µg/mL streptomycin, 350 µg/mL glutamine, and 10% inactivated fetal calf serum (FCS) (GIBCO, Carlsbad,CA). Jurkat cells were cultured in RPMI 1640 with 120 µg/mL penicillin, 120 µg/mL streptomycin, 350 µg/mL glutamine, and 10% FCS.

*CD4 T cell preparation:* CD4 T cells were isolated from fresh human peripheral blood mononuclear cells using lymphocyte separation medium (Biocoll separating solution; Biochrom, Berlin, Germany) and the RosetteSep Human CD4 +T Cell Enrichment Cocktail (Stem Cell Technologies, Vancouver, Canada). Cells were maintained in RPMI medium supplemented with l-glutamine, penicillin (50 U/ml), streptomycin (50 µg/ml), and 10% fetal bovine serum.

*Peptide synthesis:* Peptides were chemically synthesized and purified by high-performance liquid chromatography (HPLC) with a purity of >95% by Synpeptide Co.,Ltd. (Shanghai, China). EF-C has the sequence QCKIKQIINMWQ. ATTO 647N and ATTO 495 NHS ester were purchased from ATTO-TEC GmbH (Siegen, Germany, order no. ATTO 647N NHS ester: AD 647N-31 and ATTO 495 NHS ester: AD 495-31). To generate labeled fibrils,1% ATTO 647N- and ATTO 495-labeled EF-C peptides from the labeling reaction were mixed with 99% unlabeled and untreated peptide in 99.99% DMSO [1]. The fibrillation process was initiated by diluting the peptide mixture 10-fold in PBS. Fibrils were formed immediately as described previously [2]. Labeled fibrils were washed by centrifugation at 20817g for 15 min and resuspended in an equal amount of PBS.

*Scanning electron microscopy (SEM):* HeLa cells were seeded on cover glasses in 12-well plates. Jurkat T cells and freshly isolated CD4+ T cells were cultured in suspension as stated above. Adherent cells and suspension cells were incubated with EF-C PNF fibrils with a final concentration of 5 µg/ml for 1 hour at 37°C. Two different protocols were then used for SEM sample preparation. Cell media of adherent cells was removed and cells were fixed with 2.5 % glutaraldehyde (in PBS with 1% saccharose, pH7.3) over night at 4 °C. The protocol for suspension cells was derived from a previously published protocol [3]. Briefly, cells were fixed in suspension by adding 2.5ml 2.5 % glutaraldehyde (in phosphate buffer with 1% saccharose, pH7.3) to 125µl cells in medium and incubated over night at 4 °C. The next day cells were washed with PBS, resuspended and a 100 µl drop containing 1,5 x $10^5$ cells was placed onto a poly-L-lysin treated cover glass. The cells were left in a moist chamber over night at 4°C to settle onto the poly-l-lysin treated glass surface. Further preparation of adherent and suspension cells was done as described in [4]. Briefly, cells on cover glasses were post-fixed with 2% osmiumtetroxide in PBS for 20 min, washed in PBS, gradually dehydrated with increasing propanol concentrations, critical point dried and platinum coated before they were imaged using a Hitachi S-5200 field emission scanning electron microscope.

*Scanning transmission electron microscopy (STEM) tomography:* STEM tomography was performed as described previously [5,6]. For this, HeLa cells were grown on carbon-coated sapphire discs (3-mm diameter, 50-µm thickness; Wohlwend GmbH), treated for 24h with 1mg/ml EF-C PNF at 37°C before they were fixed by high-pressure freezing (HPF Compact 01, Wohlwend GmbH) and freeze substitution (EM AFS2, Leica) and embedded in Epon. 800 nm thick sections of cells were mounted on poly-L-lysine (10% in water) treated EM grids with 200 parallel grid bars (Plano). Mounted sections were again treated with poly-L-lysine to attach colloidal gold fiducials (Aurion) before they were finally coated with carbon (BAF 300, Balzers). Tomogram acquisition was conducted on a JEM-2100F (Jeol) with an accelerating voltage of 200 kV. Tilt series were acquired from -72° to +72° with a 1.5° increment. Image series were reconstructed to tomograms with the IMOD software package[7]. Three-dimensional visualization of the plasma membrane and fibrils was performed with Avizo 9 Lite (FEI/Thermo Fisher) by manual segmentation of the plasma membrane and threshold segmentation of fibrils.

*Generation of viral particles:* Lentiviral particles were generated by transfection of pBRHIV-1 NL4-3 92TH014[8] into HEK293T cells as described[9]. After transfection and overnight incubation, the transfection mixture was replaced with 2 mL of cell culture medium. Cell supernatant was harvested and centrifuged for 3 min with 13000 rpm 40h after transfection to remove cell debris.

*Generation of transduced cells:* HEK293T cells were cotransfected with pCMVdR8.91, VSV-G, pAdvantage and an empty shRNA vector containing a puromycin resistance to generate viral particles. HeLa cells stably expressing LifeAct.GFP were then transduced and 2 days post infection treated with puromycin to eradicate not transduced cells.

*Flow cytometry:* For flow cytometry, 200 000 TZM-bl cells were seeded in 12 well plates in 1ml (Sarstedt, Nümbrecht, Germany) 1 day prior to the experiment. The medium was changed to DMEM without FCS 16h before adding dynasore to prime cells. 250 000 freshly isolated CD4 T cells or Jurkat cells were seeded in 200µl RPMI. All cells were pre-incubated with indicated concentrations of dynasore for 45 min. Then 10 or 12.5 µg/ml ATTO495 labeled EF-C PNF was added to HeLa or Jurkat and CD4+ T cells for either 10, 30 or 60 minutes. Cells were washed with PBS and HeLa cells detached by pipetting. Samples were analyzed using a BD Canto II (Heidelberg, Germany) and gated for living cells and singlets.

*Confocal microscopy:* To assess cellular binding of viral particles, HeLa cells were seeded in µ-slides (8 well; Ibidi, Munich, Germany) 1 day prior to treatment. The medium was changed to DMEM without FCS 1h prior to the experiments, followed by treatment with 200 µM dynasore or DMSO for 45 min. Next, HIV-1 or a mixture of HIV-1 and EF-C PNF was added for 1h at 4 °C that a final concentration of 5 µg/ml of EF-C PNF and 1:20 diluted HIV-1 in 200µl was present on cells. Then unbound virus was removed by washing two times with PBS, followed by permeabilization and fixation of the cells by treatment with BD Cytoperm/Cytofix solution (BD Pharmingen). Finally, cells were stained with α-p24 PE (Coulter Clone KC57-RD1) and DAPI for 45 min at 4C. Samples were imaged using the Zeiss LSM710 confocal microscope. All pictures were processed equally by ImageJ using rolling ball background subtraction and adjustment of brightness and contrast. To quantify the average of virus particles per cell, virus particles were counted by two individual persons.

*Fusion Assay:* HEK293T cells were cotransfected with pNL4-3 92TH014 proviral DNA, pCMV-BlaM-Vpr, and pAdVAntage vectors as previously described[10]. After 2 days of cultivation, the virus containing supernatant was centrifuged for 10 min at 1300 rpm to remove cellular debris. Virus stocks were stored at −80 °C. Cells were pre-treated with 200µM dynasore for 45 min then incubated with virions containing BlaM-Vpr at 37°C for 2.5 h, washed in $CO_2$-independent medium (GibCo BRL, Rockville, MD), and then loaded with CCF2 dye and incubated overnight at room temperature. Next day cells were fixed with 4% paraformaldehyde and the change in emission fluorescence of CCF2 after cleavage by the BlaM-Vpr chimera was monitored with a BD LSRII. Data were analyzed with FACSDiva software (BD).

*Cell viability:* Possible effects of dynasore on the metabolic activity on HeLa cells and Jurkat cells were analyzed under the same conditions as described above. 10 000 HeLa cells or 20 000 Jurkat cells were seeded into 96-well plates and treated with indicated concentrations of dynasore for indicated time points. The Celltiter-glow assay was performed according to the manufacturer protocol. The supernatant of adherent HeLa cells was discarded and 50µl PBS as well as 50µl Celltiter-glow reagent was added to the cells for 10 min followed by measuring the luminescence. For soluble Jurkat cells, Celltiter-glow reagent was directly added to the supernatant.
For MTT, supernatant was discarded and 100 µL of 0.5 mg/mL MTT-PBS (3-[4,5-dimethyl-2-thiazolyl]-2,5- diphenyl-2H-tetrazolium bromide) solution was added to the cells. Live cells reduce the yellow MTT salt by NAD(P)H-dependent oxidoreductase system causing the formation of insoluble purple formazan crystals. After 2.5 hour, the cell-free supernatant was discarded and formazan crystals were dissolved in 100 µL dimethylsulfoxide (DMSO):Ethanol (1:1) and absorption was detected at 490 nm and corrected by the background absorption at 650 nm.

*Confocal microscopy:* For microscopy HeLa cells stably expressing LifeAct.GFP were seeded in µ-slides (8 well; Ibidi, Munich, Germany) 1 day prior to treatment. To analyze the effect of dynasore on HeLa and Jurkat cells, medium was changed to DMEM without FCS 16h prior to the experiment. Then cells were pre-incubated for 45 min with 200µM dynasore followed by direct analysis or staining with CellMask™ Deep Red Plasma membrane Stain (Invitrogen) of Jurkat cells, according to the manufacturer's protocol. All pictures were processed equally by ImageJ using rolling ball background subtraction and adjustment of brightness and contrast. For analysis of transduced HeLa cells, cells were seeded in µ-slides and analyzed the next day. For time laps movies, cells were treated with 5 µg/ml EF-F ATTO647 and directly imaged at 20 sec frame intervals over 6-20 min. All samples were imaged using the Zeiss LSM710 confocal microscope. To quantify the average of virus particles per cell, virus particles were counted by two individual persons.

**Supplementary figures:**

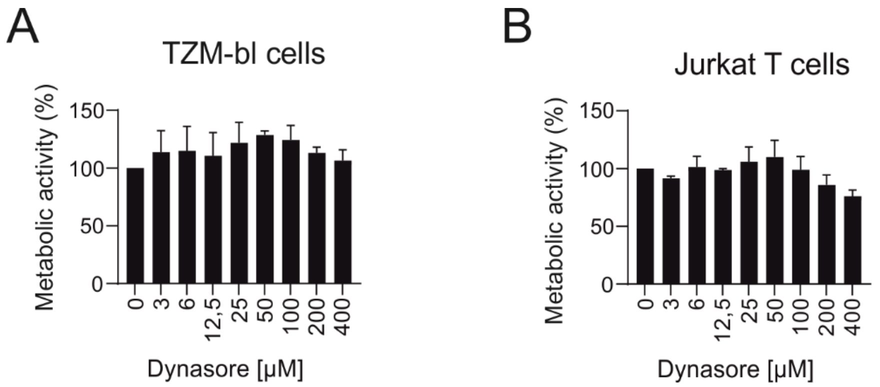


Figure S1: Cell titer glow of HeLa (A) or Jurkat cells (B) that were treated for 2h with indicated concentrations of dynasore followed by removing medium and adding CellTiter-Glo® substrate. C: MTT Assay of HeLa cells treated for 4 or 24h with indicated concentrations of dynasore. Shown are average values (± SD) of triplicate measurements from three independent experiments.

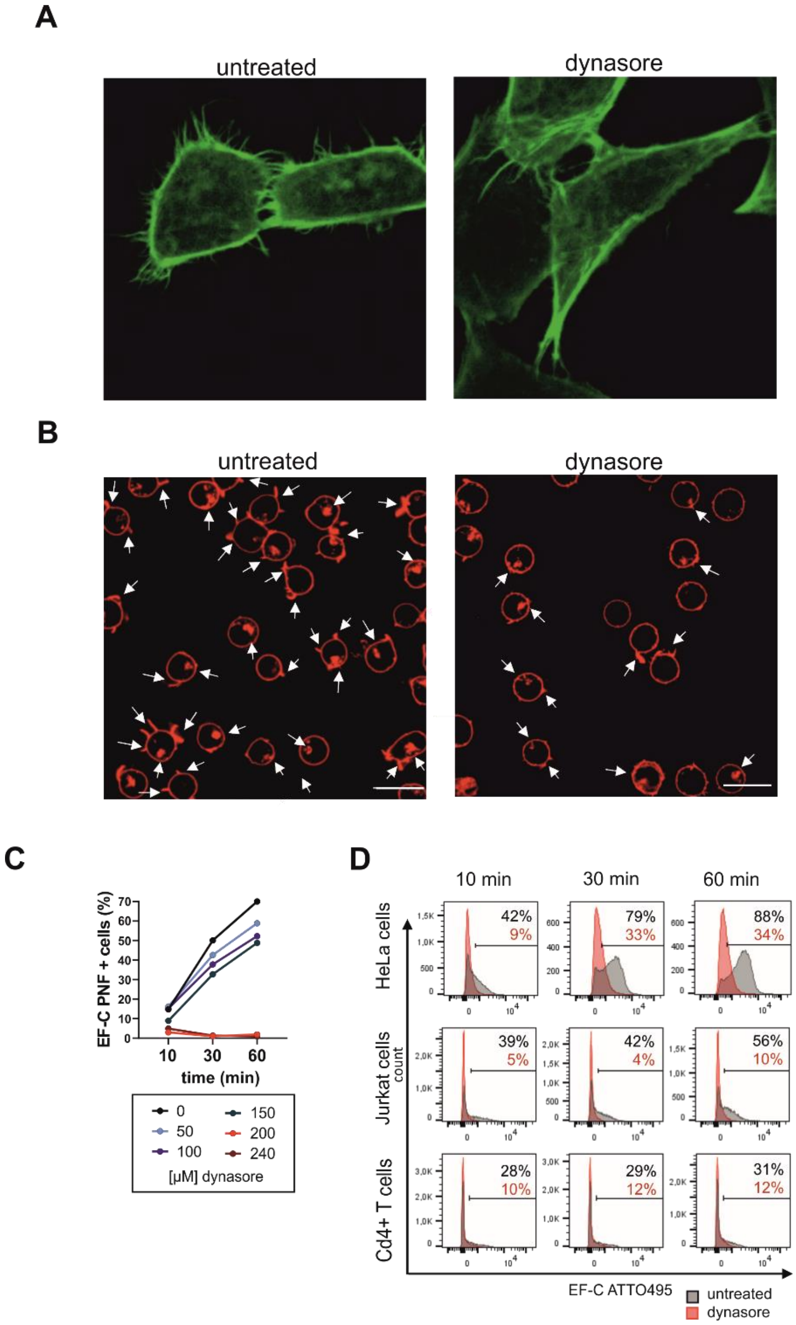


**Figure S2: Confocal microscopy analysis of HeLa (A) and Jurkat cells (B) treated for 45min with 200 µM dynasore show reduced protrusion formation.** Membrane of Jurkat cells was stained using CellMask™ Deep Red Plasma membrane Stain. Protrusions on Jurkat cells are marked with a white arrow. Scale bars are 20 µm. C: Quantification of EF-C PNF binding to HeLa cells after incubation with various concentrations of dynasore. Cells were treated with dynasore followed by adding 10mg/ml EF-C PNF for indicated time points. Shown values from one independent experiment. D: Representative FACS plots of Figure 2.

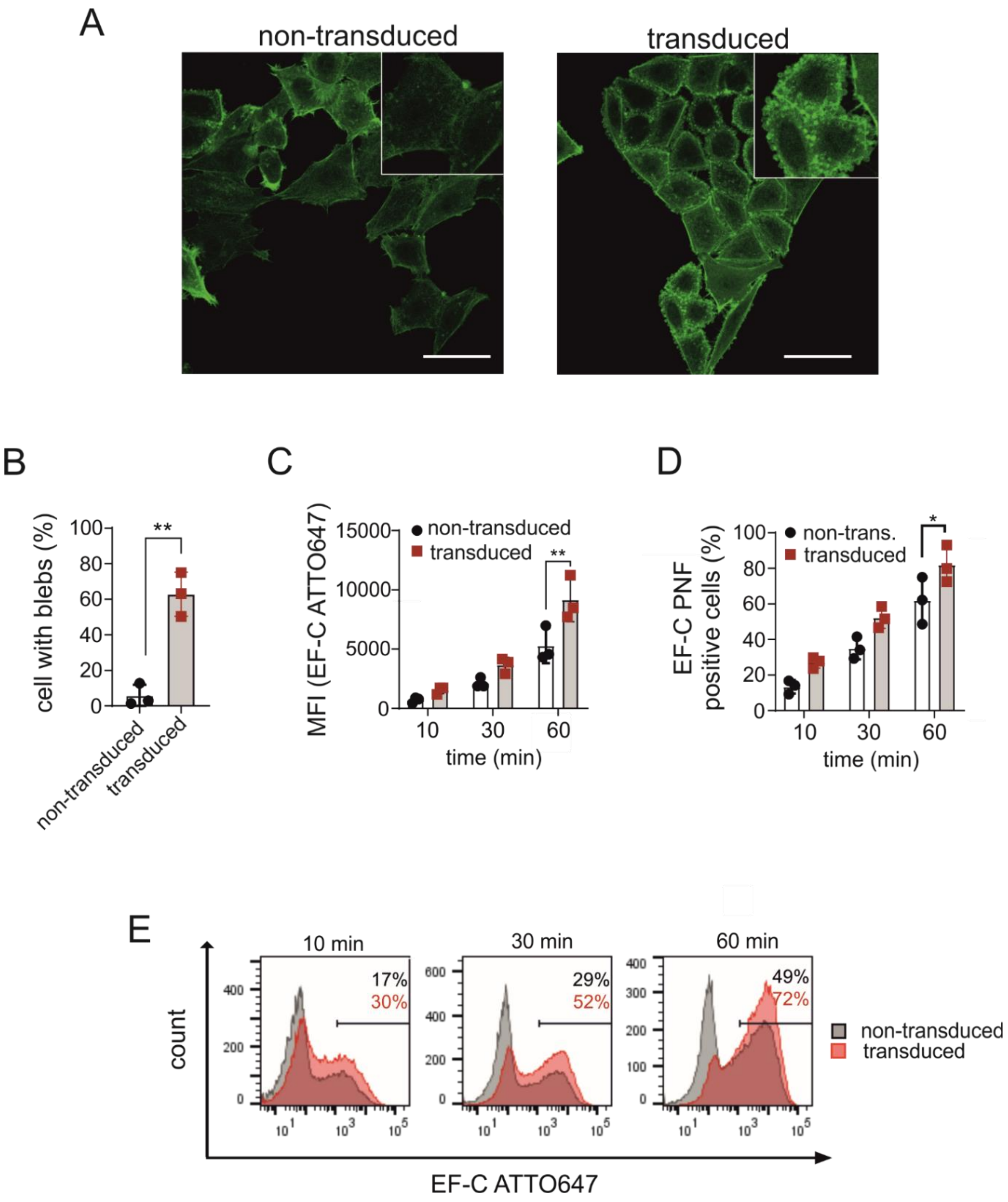


**Figure S3: Increase in protrusion through transduction lead to a increased EF-C PNF binding.** Confocal microscopy images of untreated (A) or transduced HeLa cells (B-F). Cells were transduced with a control (B), ARP3 (C), DIAPH3 (D,E) and FMNL2 (F) shRNA followed by selection by Puromycin. Cellular actin was labeled with LifeAct-GFP (green). H: Quantification of cells having blebs. Shown values (± SD) from three independent experiments. Statistical differences to untreated was identified by one-way ANOVA. I: Mean fluorescence intensity of untreated or transduced cells that were treated with 10 µg/ml ATTO647 labeled EF-C PNF. Percentage of EF-C PNF positive cells (J) and FACS plots (K) of the same experiments. Shown values (± SD) from three independent experiments. Statistical differences to untreated was identified using ordinary two-way ANOVA.

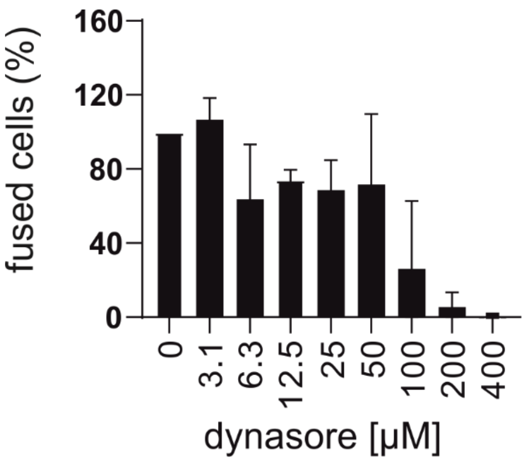


**Figure S4: Dynasore inhibits HIV-1 fusion. HIV-1 fusion was determined using BLAM Vpr reporter construct.** Cells were incubated for 45 min with 200 µM dynasore followed by adding BLAM-Vpr containing virions for 2.5h.

**Movie S1:** Time-lapse fluorescence microscopy of the interaction of EF-C ATTO647 PNF with HeLa cells. HeLa cells were exposed to 5 µg/ml EF-C ATTO647 imaged at 20 sec frame intervals over 6-20 min.

**Movie S2:** Time-lapse fluorescence microscopy of the interaction of EF-C ATTO647 PNF with HeLa GFP cells. HeLa cells stably expressing LifeAct.GFP were exposed to 5 µg/ml EF-C ATTO647 imaged at 20 sec frame intervals over 6-20 min.
